# RAD51AP1 and RAD54 underpin two distinct RAD51-dependent routes of DNA damage repair via homologous recombination

**DOI:** 10.1101/2021.07.15.452469

**Authors:** Platon Selemenakis, Neelam Sharma, Youngho Kwon, Mollie Uhrig, Patrick Sung, Claudia Wiese

## Abstract

Homologous recombination (HR) is a complex DNA damage repair pathway and an attractive target of inhibition in anti-cancer therapy. To help guide the development of efficient HR inhibitors, it is critical to identify compensatory sub-pathways.

In this study, we describe a novel synthetic interaction between RAD51AP1 and RAD54, two structurally unrelated proteins that function downstream of the RAD51 recombinase in HR. We show that deletion of both RAD51AP1 and RAD54 synergistically sensitizes human cancer cell lines to treatment with a Poly(adenosine 5’
s-diphosphate-ribose) polymerase inhibitor, to the DNA inter-strand crosslinking agent mitomycin C, and to hydroxyurea, which stalls the progression of DNA replication forks. We infer that HR-directed anti-cancer treatment modalities shall consider this intra-pathway functional overlap, and we hypothesize that in cancerous cells the simultaneous inactivation of both RAD54 and RAD51AP1 will accentuate tumor kill.

## Introduction

Homologous recombination (HR) is an essential DNA damage repair pathway critical for genome stability and tumor suppression. HR is altered in many different tumor types and has become an attractive target for the development of new anti-cancer therapies (Kopa et al. 2019; Trenner and Sartori 2019). Accurate HR is restricted to the S- and G2-phases of the cell cycle where the sister chromatid is used as the template for the restoration of lost sequence information at the damaged DNA site. At the DNA break, a 3’
s-single-stranded (ss)DNA overhang is generated and protected by the ssDNA-binding protein RPA (Symington 2014; Daley et al. 2015). RPA is replaced by the RAD51 recombinase, a rate-limiting step in the HR reaction that is facilitated by multiple recombination mediators (Sung 1997a; Sung 1997b; Dosanjh et al. 1998; Sung et al. 2003; Zhao et al. 2015; Belan et al. 2021; Roy et al. 2021). The RAD51-ssDNA nucleoprotein filament catalyzes the capture of the DNA template and initiates the formation of a displacement loop (D-loop) with the assistance of several RAD51-associated proteins (Petukhova et al. 1998; Tanaka et al. 2000; Miyagawa et al. 2002; Modesti et al. 2007; Wiese et al. 2007; Zhao et al. 2017).

RAD51AP1 and RAD54 are two RAD51-associated proteins that co-operate with the RAD51 filament in the capture of the DNA donor molecule and in formation of the D-loop (Petukhova et al. 1998; Tanaka et al. 2000; Miyagawa et al. 2002; Modesti et al. 2007; Wiese et al. 2007; Zhao et al. 2017). RAD51AP1 may have evolved in response to the higher complexities of vertebrate genomes (Parplys et al. 2014). In contrast, RAD54 is highly conserved across eukaryotes (Clever et al. 1997; Essers et al. 1997; Golub et al. 1997; Petukhova et al. 1998; Swagemakers et al. 1998). RAD51AP1 functions in the protection of cells from genotoxic agents, in genome stability, in the HR-mediated alternative lengthening of telomeres (ALT) pathway and promotes HR when local transcription is active (Henson et al. 2006; Modesti et al. 2007; Wiese et al. 2007; Barroso-Gonzalez et al. 2019; Ouyang et al. 2021). Similarly, RAD54 maintains HR capability, cell survival after treatment with chemotherapeutic agents, and ALT activity (Swagemakers et al. 1998; Tan et al. 1999; Mason et al. 2015; Spies et al. 2016; Mason-Osann et al. 2020). Strikingly, in human cells, loss of either RAD51AP1 or RAD54 engenders only mild HR-deficiency (Henson et al. 2006; Modesti et al. 2007; Wiese et al. 2007; Gottipati et al. 2010; Spies et al. 2016; Olivieri et al. 2020).

The RAD51AP1 and RAD54 proteins are unrelated structurally, but both upregulate RAD51 activity by enhancing the ability of the RAD51 filament to engage with the homologous double-stranded (ds)DNA donor (*i*.*e*., in synapsis) and in strand invasion (Petukhova et al. 1998; Petukhova et al. 1999; Solinger and Heyer 2001; Solinger et al. 2001; Sigurdsson et al. 2002; Modesti et al. 2007; Wiese et al. 2007). In these steps of the HR reaction, RAD51AP1 may serve as an anchor between the two DNA molecules undergoing exchange (Modesti et al. 2007; Dunlop et al. 2012; Pires et al. 2021). In contrast, RAD54 belongs to the SWI2/SNF2 protein family of DNA-dependent ATPases (Flaus et al. 2006) and utilizes its ATPase activity to convert the synaptic complex into a D-loop (Sigurdsson et al. 2002; Crickard et al. 2020), and to translocate along the DNA (Van Komen et al. 2000; Ristic et al. 2001) whereby chromatin is remodelled and the turnover of RAD51 is facilitated (Alexiadis and Kadonaga 2002; Alexeev et al. 2003; Jaskelioff et al. 2003; Li and Heyer 2009).

The mild phenotype of RAD54-deficient human cells has been attributed to the existence of RAD54B, a RAD54 paralog (Hiramoto et al. 1999). Human RAD54 and RAD54B share 48% identity and 63% similarity (Flaus et al. 2006; Ceballos and Heyer 2011). Although less well understood than RAD54, existing evidence implicates RAD54B in the core mechanisms of HR (Tanaka et al. 2000; Miyagawa et al. 2002; Flaus et al. 2006; McManus et al. 2009; Ceballos and Heyer 2011). Compared to RAD54, RAD54B was identified as the weaker ATPase, and these results suggest that RAD54B may fulfil a backup role for RAD54 (Tanaka et al. 2002).

In this study, we show that loss of *RAD54* in human cells can be compensated for by the RAD51AP1 protein. We show that simultaneous deletion of the *RAD54* and *RAD51AP1* genes synergistically sensitizes human cancer cell lines to treatment with the DNA inter-strand crosslinking agent mitomycin C (MMC), prolonged exposure to replication stalling by hydroxyurea (HU), and to Poly(adenosine 5’
s-diphosphate-ribose) polymerase inhibition (PARPi). We also show that the RAD54 paralog RAD54B can substitute for RAD54 activity, but, surprisingly, to a lesser degree than RAD51AP1. Based on these results, we conclude that the activities of RAD51AP1 and RAD54 underpin two major, mechanistically distinct routes for the completion of HR in human cells.

## Results

### Deletion of both RAD54 and RAD51AP1 synergistically sensitizes human cancer cell lines to the cytotoxic effects of MMC exposure

To investigate the genetic interaction between RAD51AP1 and RAD54, we generated *RAD54*/*RAD51AP1* double knockout (KO) HeLa cell lines and compared the phenotypes of these double KO cells to HeLa cells deleted for either *RAD51AP1* or *RAD54* (Liang et al. 2019; Maranon et al. 2020). To generate *RAD54*/*RAD51AP1* double KO cells we targeted *RAD54* by CRISPR/Cas9-nic in *RAD51AP1* KO cells and selected two of several *RAD54*/*RAD51AP1* double KO clones for the experiments described below. We verified the loss of protein expression by Western blot analysis (Fig. 1A, lanes 5-6), sequenced across the Cas9-nic cleavage sites in *RAD54* to confirm mutagenesis (Supplemental Fig. S1A-C), and used immunocytochemistry (ICC) to monitor the loss of RAD54 foci formation after γ-irradiation (Supplemental Fig. S1D).

**Figure 1.**
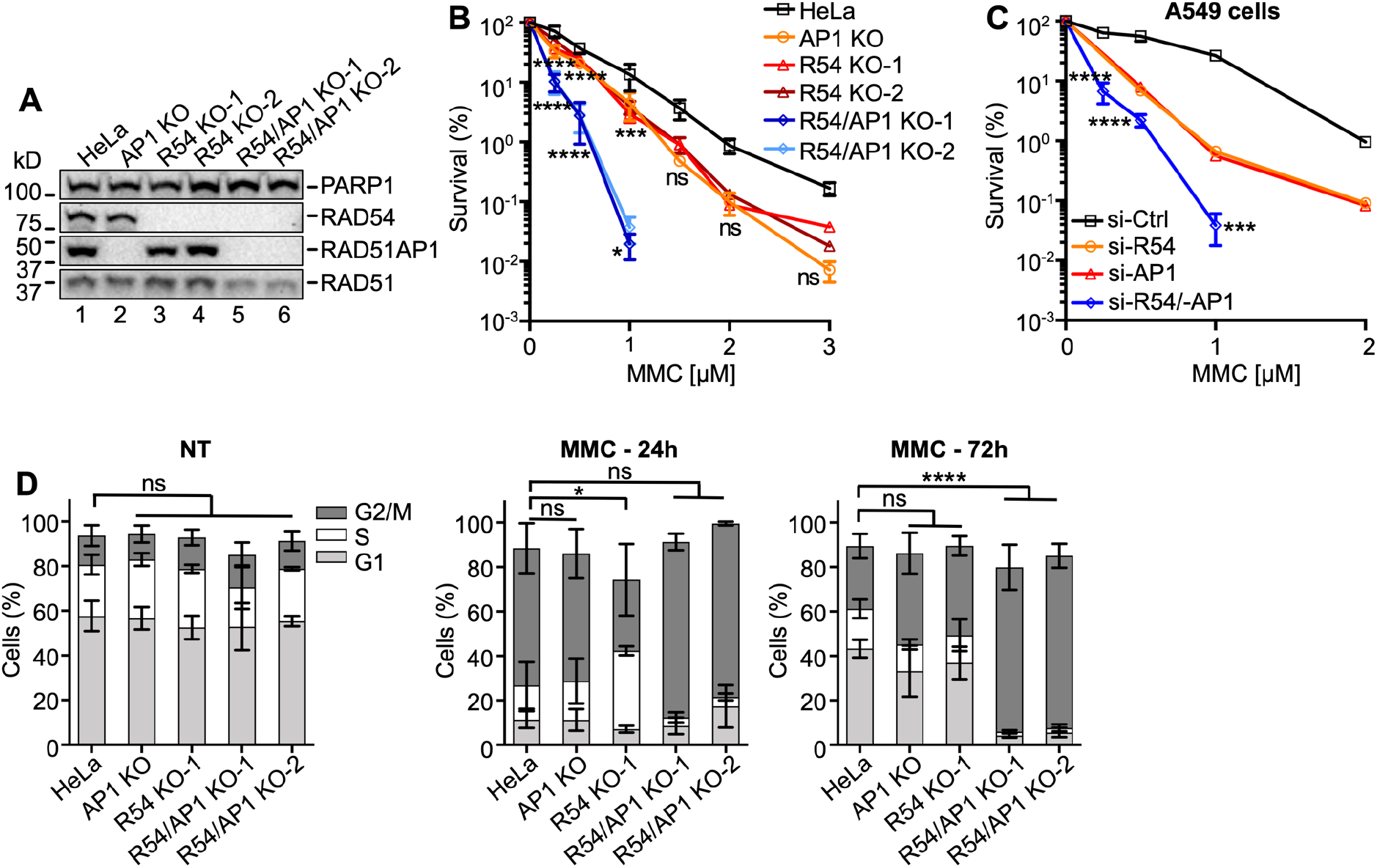
Deficiency of both *RAD51AP1* and *RAD54* synergistically affects MMC cytotoxicity and cell cycle progression. (*A*) Western blots of nuclear extracts of HeLa cells and derivatives. *RAD51AP1* KO cells (here: AP1 KO), two independently isolated *RAD54* KO cell lines (here: R54 KO-1, R54 KO-2), and two independently isolated *RAD54*/*RAD51AP1* KO cell lines (here: R54/AP1 KO-1; R54/AP1 KO-2). The signal for PARP1 serves as loading control. (*B*) Results from clonogenic cell survival assays after MMC. Data points are the means from 2-5 independent experiments ±SD. ***, *p* < 0.001; ****, *p* < 0.0001; ns, non-significant; two-way ANOVA test followed by Dunnett’s multiple comparisons test. (*C*) Results from clonogenic cell survival assays of MMC-treated A549 cells with RAD51AP1 and/or RAD54 knockdown. Data points are the means from three technical replicates for A549 cells transfected with RAD54 siRNA (here: si-R54) or RAD51AP1 siRNA (si-AP1), and from two independent experiments ±SD for A549 cells transfected with negative control siRNA (si-Ctrl) or RAD54 and RAD51AP1 siRNA (si-R54/-AP1). ***, *p* < 0.001; ****, *p* < 0.0001; two-way ANOVA test followed by Sidak’s multiple comparisons test. (*D*) Average percentage of cells in G1, S and G2/M cell cycle phases without (here: NT (no treatment)), and 24 and 72 h after release from MMC. Bars are the means from at least three independent experiments ±SD. *, *p* < 0.05, ****, *p* < 0.0001, ns, non-significant; one-way ANOVA test followed by Dunnett’s multiple comparisons test.

We determined the growth rates of all HeLa cell derivatives (*i*.*e*., single KO and double KO cells) and detected no significant differences in population doubling times (Supplemental Fig. S1E). In fractionated protein extracts from unperturbed cells, we noted higher levels of RAD54 protein in *RAD51AP1* KO cells (Supplemental Fig. S1F, lanes 2 and 8) and higher levels of RAD51AP1 protein in *RAD54* KO cells (Supplemental Fig. S1F, lanes 9-10).

Next, we tested the sensitivity to MMC of single and double KO cells in clonogenic cell survival assays. In accord with our earlier studies (Liang et al. 2019; Maranon et al. 2020), we show that *RAD51AP1* and *RAD54* single KO cells are moderately sensitized to the cytotoxic effects of MMC (Fig. 1B). Deletion of both *RAD51AP1* and *RAD54*, however, synergistically sensitized HeLa cells to MMC (Fig. 1B), suggesting a non-epistatic relation between RAD51AP1 and RAD54. To exclude that this effect was specific to HeLa cells, we depleted RAD51AP1 and/or RAD54 in A549 lung cancer cells (Supplemental Fig. S1G). A549 cells depleted for either RAD51AP1 or RAD54 showed similarly increased sensitivities to MMC while loss of both RAD51AP1 and RAD54 rendered A549 cells significantly more sensitive to the cytotoxic effects of MMC (Fig. 1C). Collectively, these results reveal compensation between RAD51AP1 and RAD54 in the protection of human cells from MMC-induced DNA damage.

We used U2OS-DRGFP cells (Nakanishi et al. 2005; Xia et al. 2006) to assess the effects of RAD51AP1 and/or RAD54 depletion on gene conversion. Depletion of both RAD51AP1 and RAD54 downregulated the levels of gene conversion at DRGFP ∼ 10-fold (*p* < 0.001; Supplemental Fig. S1H), while single knockdown of either RAD51AP1 or RAD54 impaired gene conversion ∼ 2-fold, as previously shown (Wiese et al. 2007; Spies et al. 2016).

Next, we assessed cell cycle progression upon MMC exposure of single and *RAD54*/*RAD51AP1* double KO cells and compared the results to HeLa cells. In the absence of MMC, all cell lines progressed similarly through the cell cycle (Fig. 1D, left panel; Supplemental Fig. S1I). Twenty-four hours after release from MMC, all cell lines remained arrested in cell cycle progression (Fig. 1D, middle panel; Supplemental Fig. S1I). At 72 h post release from MMC, HeLa cells, *RAD51AP1* KO, and *RAD54* KO cells regained the capacity to proceed through mitosis and enter the following cell cycle. Both *RAD54/RAD51AP1* double KO cell lines, however, remained arrested in G2/M phase (*p* < 0.0001; Fig. 1D, right panel; Supplemental Fig. S1I), likely due to their higher fraction of unresolved or mis-repaired DNA damage.

### RAD54 deficiency is rescued by ectopic RAD54

Ectopic expression of HA-tagged RAD54 in *RAD54*/*RAD51AP1* double KO cells reverted their response to MMC to the level of *RAD51AP1* KO cells (Fig. 2A; Supplemental Fig. S2A). Similarly, cell cycle progression after MMC of *RAD54*/*RAD51AP1* double KO cells with ectopic RAD54 became similar to that of *RAD51AP1* KO cells (Fig. 2B; Supplemental Fig. S1I). Moreover, *RAD54*/*RAD51AP1* double KO cells with ectopic RAD54 formed RAD54 foci after γ-irradiation (Supplemental Fig. S1D). Ectopic expression of RAD54 also rescued the sensitivity to MMC of single *RAD54* KO cells (Supplemental Fig. S2B,C) and RAD54 foci formation after γ-irradiation (Supplemental Fig. S1D). These results show that the phenotypes associated with *RAD54* deficiency in *RAD54*/*RAD51AP1* double KO and *RAD54* single KO cells stem from the loss of *RAD54*. However, ectopic expression of high amounts of RAD54 in *RAD51AP1* KO cells (Supplemental Fig. S2A, lanes 3-4) did not rescue their sensitivity to MMC (Fig. 2C), demonstrating that defined attributes of the RAD51AP1 protein cannot be compensated for by RAD54.

**Figure 2.**
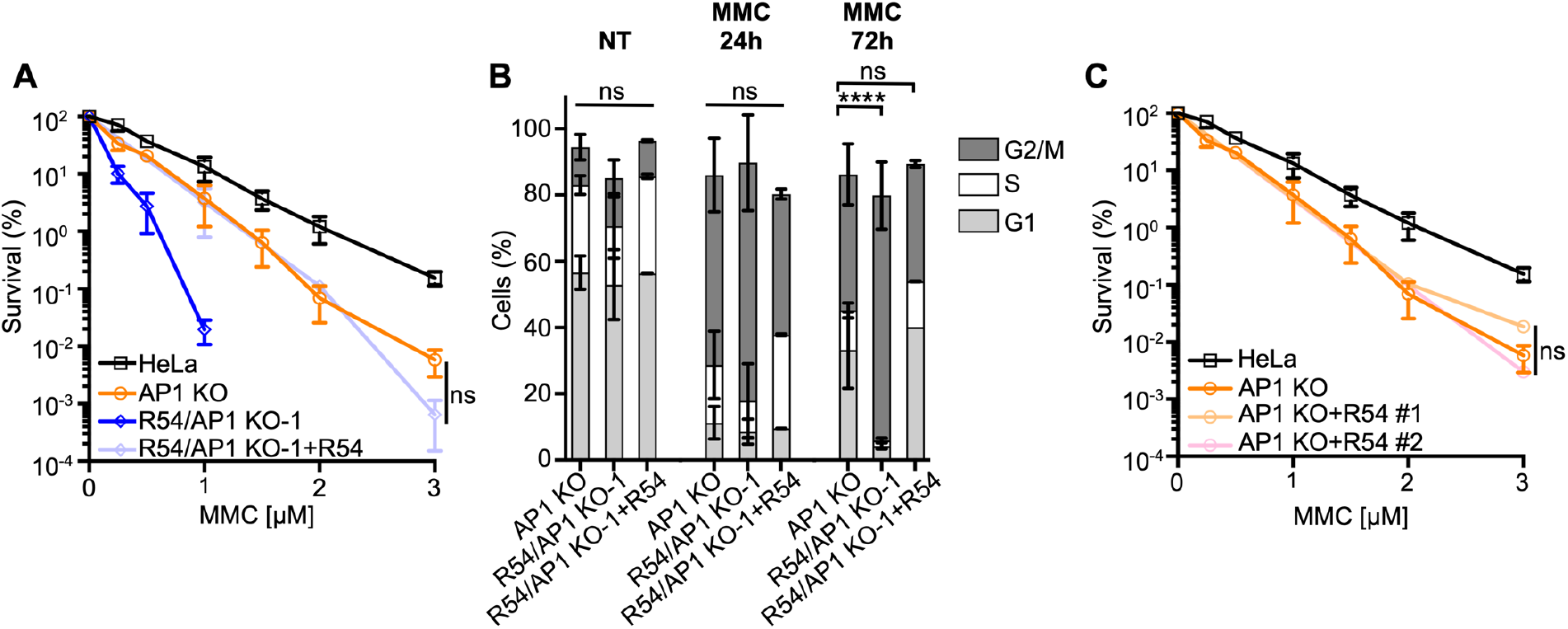
Ectopic expression of RAD54 rescues *RAD54* deficiency in *RAD54*/*RAD51AP1* double KO cells. (*A*) Results from MMC clonogenic cell survival assays of *RAD54*/*RAD51AP1* double KO cells (R54/AP1 KO-1) with (here: +R54) and without ectopic RAD54 and of *RAD51AP1* KO (AP1 KO) and HeLa cells for comparison purposes. Data points are the means from two independent experiments ±SD. ns: non-significant; two-way ANOVA test followed by Tukey’s multiple comparisons test. (*B*) Average percentage *RAD54*/*RAD51AP1* double KO cells with (here: +R54) and without ectopic RAD54 and of *RAD51AP1* KO cells in G1, S and G2/M cell cycle phases without MMC (NT), and 24 h and 72 h after release from MMC. Bars represent the means from two independent experiments ±SD. ****, *p* < 0.0001; ns, non-significant; one-way ANOVA test followed by Dunnett’s multiple comparisons test. (*C*) Results from MMC clonogenic cell survival assays of *RAD51AP1* KO cells and two independently isolated *RAD51AP1* KO clones expressing different amounts of ectopic RAD54-HA (see Supplemental Fig. S2A). Data points are the means from two independent experiments ±SD for AP1 KO+R54 #1 cells and from three technical replicates for AP1 KO+R54 #2 cells. ns, non-significant; two-way ANOVA test followed by Tukey’s multiple comparisons test.

### Deletion of both RAD51AP1 and RAD54 synergistically sensitizes HeLa cells to replication stress

We treated all cell lines with 4 mM HU for 5 hours, which blocks DNA synthesis and stalls replication fork movement (Liu et al. 2020). To understand the fate of stalled replication forks in single and double KO cells, we monitored the recovery of cells from stalled replication using the single-molecule DNA fiber assay. We pulse-labeled cells with the thymidine analog 5-Chloro-2’
s-deoxyuridine (CldU) first, then replenished cells with HU-containing medium prior to pulse-labeling with 5-Iodo-2’-deoxyuridine (IdU) (Fig. 3A). We then determined the ability of all cell lines to restart DNA replication by measuring the lengths of IdU tracts of CldU-labeled DNA fibers (Fig. 3B; Supplemental Fig. S3D; Table S1). *RAD51AP1* KO cells showed no significant defect in fork restart compared to HeLa cells (*p* = 0.0758). In contrast, *RAD54* KO cells restarted forks significantly faster than HeLa cells (*p* < 0.01), possibly related to the role of RAD54 in catalyzing fork regression (Bugreev et al. 2011). However, in comparison to both HeLa and single KO cells, *RAD54*/*RAD51AP1* double KO cells were significantly impaired in fork restart (*p* < 0.0001; Fig. 3C; Supplemental Fig. S3D; Table S1). Collectively, these results show that the efficient restart from stalled replication relies on RAD51AP1 or RAD54 in HeLa cells. The results also suggest that RAD54 suppresses accelerated fork restart after HU.

**Figure 3.**
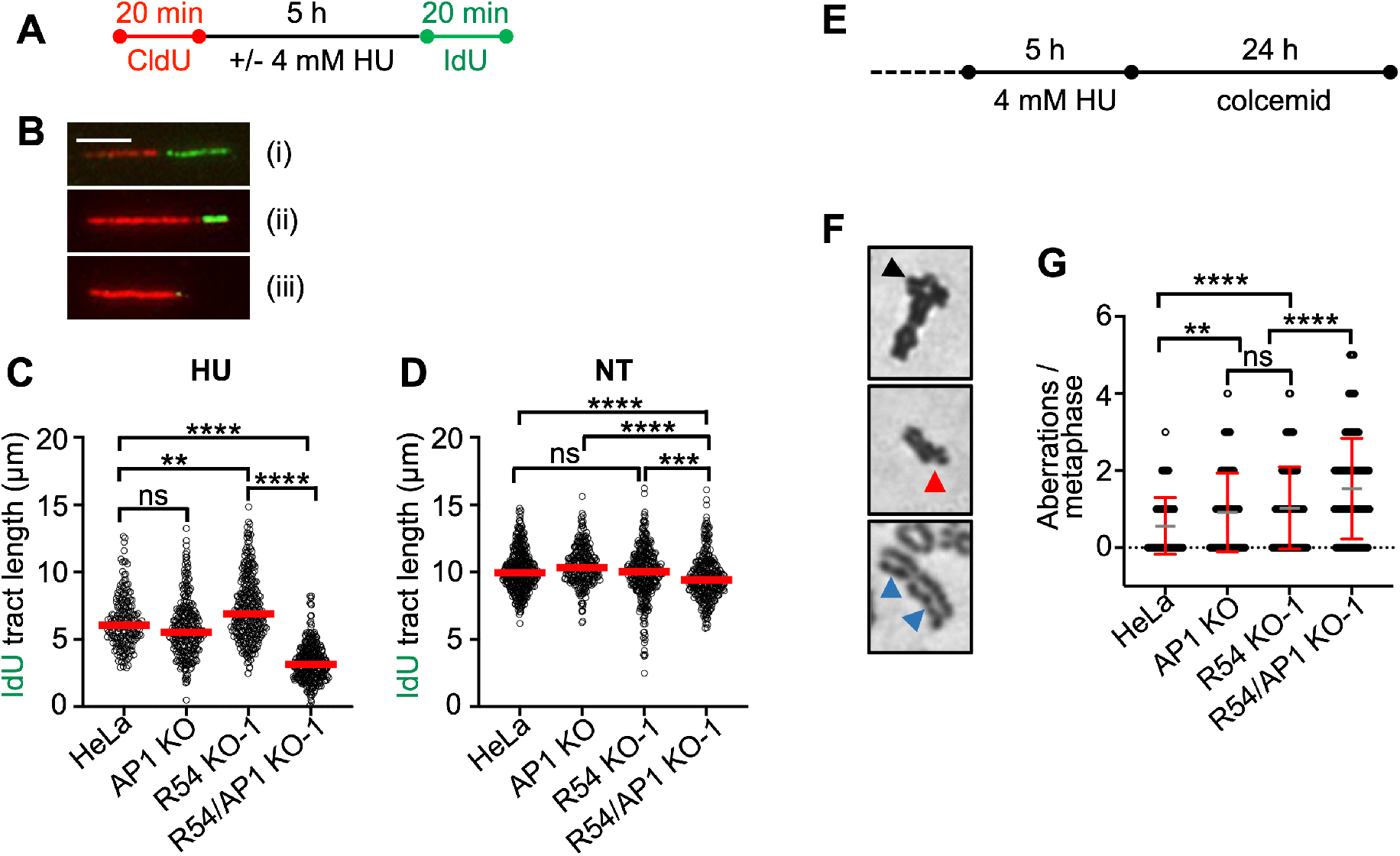
Concomitant loss of *RAD51AP1* and *RAD54* results in increased replication stress and genome instability. (*A*) Schematic of the experimental protocol for the DNA fiber assay. (*B*) Representative micrographs of fiber tracts of a restarted (i), a delayed (ii), and a replication fork that failed to restart (iii). Scale bar: 10 μm. (*C*) Median IdU tract length (green) after HU of *RAD51AP1* KO (AP1 KO), *RAD54* KO (R54 KO-1), and *RAD54*/*RAD51AP1* double KO (R54/AP1 KO-1) cells. Data points are from 100-150 fibers of two independent experiments each, with medians (red lines). **, *p* < 0.01; ****, *p* < 0.0001; ns, non-significant; Kruskal-Wallis test followed by Dunn’s multiple comparisons test. (*D*) Median IdU tract length (green) under unperturbed conditions (NT). Data points are from 100-150 fibers of two independent experiments each, with medians (red lines). **, *p* < 0.01; ***, *p* < 0.001; ****, *p* < 0.0001; ns, non-significant; Kruskal-Wallis test followed by Dunn’s multiple comparisons test. (*E*) Schematic of the experimental protocol to induce chromosomal aberrations. (*F*) Representative micrographs of chromosomal aberrations after HU; radial (black arrow), chromatid break (red arrow) and chromatid gaps (blue arrows). (*G*) Aberrations per metaphase after HU in *RAD51AP1* KO (AP1 KO), *RAD54* KO (R54 KO-1), and *RAD54*/*RAD51AP1* double KO (R54/AP1 KO-1) cells. Data points are from 100 metaphases of two independent experiments each. Means (grey) ±SD (red) are shown. **, *p* < 0.01; ****, *p* < 0.0001; ns, non-significant; one-way ANOVA test followed by Tukey’s multiple comparisons test.

In unperturbed cells, DNA replication progressed significantly slower in *RAD54*/*RAD51AP1* double KO cells than in HeLa or single KO cells, suggesting that endogenous obstacles to fork progression impede DNA replication in *RAD54*/*RAD51AP1* double KO cells (*p* < 0.0001and *p* < 0.001; Fig. 3D; Supplemental Fig. S3D; Table S1).

In response to replication stress, replication forks reverse into four-way junctions through annealing of the nascent DNA strands (Zellweger et al. 2015). Fork reversal is mediated by RAD51 and several DNA motor proteins and serves to bypass obstacles encountered by the replisome (Thangavel et al. 2015; Zellweger et al. 2015; Mijic et al. 2017; Thakar and Moldovan 2021). Reversed forks must be protected from nucleolytic attack to prevent fork attrition (Petermann et al. 2010; Schlacher et al. 2011; Thangavel et al. 2015; Lemacon et al. 2017; Taglialatela et al. 2017). To assess if RAD54 and/or RAD51AP1 function in the protection of replication forks from unprogrammed nuclease degradation, CldU tracts in cells exposed to HU were measured and compared to the CldU tract lengths in untreated cells (Supplemental Fig. S3A). CldU tracts after HU were shorter than those in unperturbed cells for all cell lines tested (Supplemental Fig. S3B,C; Table S1). Overall, however, CldU tracts in HU-treated *RAD51AP1* KO, *RAD54* KO, and *RAD54*/*RAD51AP1* double KO cells were not shorter than those in HU-treated HeLa cells (Supplemental Fig. S3C; Table S1). These results suggest that RAD54 and RAD51AP1 largely function independently of the protection mechanism of reversed forks in overcoming replication stress in HeLa cells. We infer that replication forks in HeLa cells and the KO cell lines are degraded as part of the normal cellular physiology in response to prolonged fork stalling by HU (Thangavel et al. 2015), and that the recruitment of proteins involved in the protection of nascent DNA at replication forks likely proceeds normally in *RAD54*/*RAD51AP1* single and double KO cells.

### Concomitant loss of RAD51AP1 and RAD54 exacerbates genome instability

Next, we tested the consequences of induced replication stress to cells with impaired replication restart. To this end, we determined chromatid gaps and breaks, and complex chromosome aberrations (*i*.*e*., radials) in HeLa, single KO and *RAD54*/*RAD51AP1* double KO cells after treatment with HU (Fig. 3E,F). Exposure to HU led to 0.56±0.73 mean aberrations per metaphase in HeLa cells and to a significant increase in mean aberrations per metaphase in both *RAD51AP1* (0.91±1.02; *p* < 0.01) and *RAD54* single KO cells (1.03±1.07; *p* < 0.0001; Fig. 3G). As expected, in *RAD54*/*RAD51AP1* double KO cells, the mean number of aberrations per metaphase was further increased compared to the single KO cells (1.53±1.30; *p* < 0.0001). These results show that replication stress leads to genome instability most prominently in *RAD54*/*RAD51AP1* double KO cells, which also show the most pronounced defect in fork restart.

### RAD54B compensates for RAD54 activity in the presence of RAD51AP1

In human cells, the role of RAD54B in HR is not well understood. In mice, however, the contribution of Rad54B to HR was discovered in the absence of Rad54 (Wesoly et al. 2006). To investigate the impact of RAD54B on the protection of human cells from MMC-induced DNA damage, we depleted RAD54B in HeLa, single KO, and *RAD54*/*RAD51AP1* double KO cells (Supplemental Fig. S4A) and performed MMC cell survival assays. Depletion of RAD54B in HeLa and *RAD51AP1* KO cells had no effect on their sensitivity to MMC (Fig. 4A). Similarly, depletion of RAD54B in *RAD54*/*RAD51AP1* double KO cells did not increase MMC cytotoxicity. In contrast, depletion of RAD54B in *RAD54* KO cells further sensitized *RAD54* KO cells to MMC (*p* = 0.044; Fig. 4A). These results show that the activity of RAD54B is critical for the protection of human cells from MMC cytotoxicity in the absence of RAD54. In HeLa, *RAD51AP1* KO, and *RAD54*/*RAD51AP1* double KO cells, however, RAD54B appears to play no detectable role in the protection of cells from MMC-induced DNA damage.

**Figure 4.**
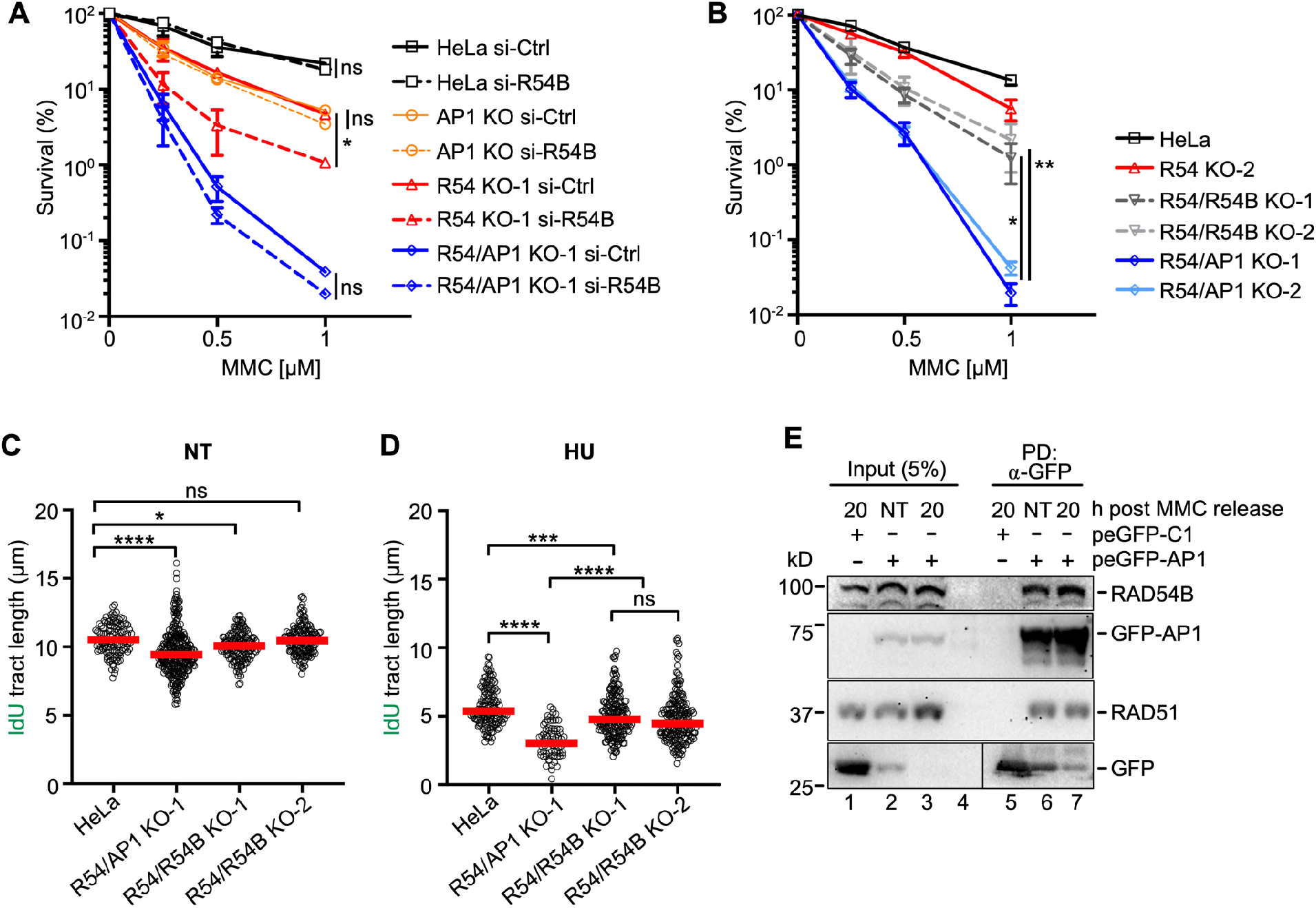
Concomitant loss of *RAD54* and *RAD54B* exacerbates MMC cytotoxicity and replication stress less extensively than concomitant loss of *RAD54* and *RAD51AP1*. (*A*) Results from clonogenic cell survival assays after MMC of cells transfected with negative control (si-Ctrl) or RAD54B siRNA (si-R54B); HeLa, *RAD51AP1* KO (AP1 KO), *RAD54* KO (R54 KO-1), *RAD54*/*RAD51AP1* double KO (R54/AP1 KO-1) cells. Data points are the means from two independent experiments ±SD. *, *p* < 0.05; ns, non-significant; two-way ANOVA test followed by Tukey’s multiple comparisons test. (*B*) Results from clonogenic cell survival assays after MMC treatment of HeLa, *RAD54* KO (R54 KO-2), *RAD54*/*RAD54B* double KO (R54/54B KO-1, KO-2), and *RAD54*/*RAD51AP1* double KO (R54/AP1 KO-1, KO-2) cells. Data points are the means from two independent experiments ±SD. *, *p* < 0.05; **, *p* < 0.01; two-way ANOVA test followed by Dunnett’s multiple comparisons test. (*C*) Median IdU tract length under unperturbed conditions (NT) of HeLa, *RAD54*/*RAD51AP1* double KO (R54/AP1 KO-1), and *RAD54*/*RAD54B* double KO (R54/54B KO-1, KO-2) cells. Data points are from 150-300 fibers of one experiment, with medians (red lines). *, *p* < 0.05; ****, *p* < 0.0001; ns, non-significant; Kruskal-Wallis test followed by Dunn’s multiple comparisons test. (*D*) Median IdU tract lengths after HU of HeLa, *RAD54*/*RAD51AP1* double KO (R54/AP1 KO-1), and *RAD54*/*RAD54B* double KO (R54/54B KO-1, KO-2) cells. Data points are from 150-200 fibers of one experiment, with medians (red lines). ***, *p* < 0.001; ****, *p* < 0.0001; ns, non-significant; Kruskal-Wallis test followed by Dunn’s multiple comparisons test. (*E*) Western blots to show that endogenous RAD54B co-precipitates in anti-eGFP protein complexes from *RAD54*/*RAD51AP1* KO cells ectopically expressing eGFP-RAD51AP1 (here: peGFP-AP1) in the absence of MMC (NT; lane 6) and 20 h after a 2-h incubation in 0.5 μM MMC (lane 7). RAD51: positive control for interaction, as previously shown in different cell types (Kovalenko et al. 1997; Wiese et al. 2007). Lane 5: Neither RAD54B nor RAD51 co-precipitate in anti-eGFP protein complexes generated from *RAD54*/*RAD51AP1* KO cells transfected with control plasmid (peGFP-C1).

To exclude the possibility that the mild increase in MMC sensitivity of *RAD54* KO cells depleted for RAD54B was the result of incomplete RAD54B knockdown, we generated *RAD54*/*RAD54B* double KO HeLa cells (Supplemental Fig. S4B-D) and compared their response to MMC to that of the *RAD54*/*RAD51AP1* double KOs. Similar to what we observed after RAD54B knockdown, two independently isolated *RAD54*/*RAD54B* double KO cells lines were significantly more resistant to MMC than *RAD54*/*RAD51AP1* double KO cells (*p* = 0.037 and p = 0.007 for *RAD54*/*RAD54B* KO-1 and KO-2, respectively; Fig. 4B). These results show that in the absence of RAD54, human cells more heavily rely on RAD51AP1 than on RAD54B to resist MMC cytotoxicity.

To understand the consequences of concomitant RAD54 and RAD54B loss on replication fork dynamics, we used the DNA fiber assay, as described above (for schematic of the protocol see Fig. 3A). As shown in Fig. 3D and herein determined independently, replication progressed significantly more slowly in *RAD54*/*RAD51AP1* double KO cells than in HeLa cells under unperturbed conditions (p < 0.0001; Fig. 4C; Table S1). In contrast, fork progression in unperturbed *RAD54*/*RAD54B* KO-1 and KO-2 cells was more similar to that of HeLa cells (Fig. 4C; Supplemental Fig. S3D; Table S1). After HU, fork restart was significantly slower in *RAD54*/*RAD51AP1* double KO cells than in *RAD54*/*RAD54B* double KO cells (*p* < 0.0001; Fig. 4D; Supplemental Fig. S3D; Table S1). These results show that, in response to stalled DNA replication in the absence of RAD54, the activities of both RAD51AP1 and RAD54B are important to efficiently restart replication forks. However, the absence of RAD51AP1 is more detrimental to the recovery from stalled replication than that of RAD54B.

### RAD54B co-precipitates in RAD51AP1 complexes

Since RAD54B knockdown did not further increase the sensitivity to MMC of *RAD51AP1* single and *RAD54/RAD51AP1* double KO cells, we hypothesized that this – in part – could be the result of RAD54B and RAD51AP1 acting in unity during the protection of cells from MMC-induced cytotoxicity. As such, we asked if RAD51AP1 may function in conjunction with RAD54B in human cells, and if a complex between these two proteins could be identified. Using the purified proteins, we previously showed that RAD51AP1 and RAD54 physically can interact, and that both proteins compete in binding to RAD51 (Maranon et al. 2020). Based on these results, we first tested if endogenous RAD54 would co-precipitate in anti-RAD51AP1 complexes of *RAD51AP1* KO cells stably expressing FLAG-tagged RAD51AP1. Our results show that RAD54 co-precipitates with FLAG-RAD51AP1 under unperturbed conditions (Supplemental Fig. S4E, lane 4).

Next, we tested the association between RAD51AP1 and RAD54B in human cells. As RAD54B activity is more prevalent in the absence of RAD54 (Fig. 4A,B), we used *RAD54*/*RAD51AP1* double KO cells with transiently expressed eGFP-tagged RAD51AP1. Both RAD51 and RAD54B were present in anti-eGFP precipitates from *RAD54*/*RAD51AP1* double KO cells expressing eGFP-RAD51AP1 (Supplemental Fig. S4F, lane 7); in contrast, RAD54B was absent in anti-eGFP precipitates from *RAD54*/*RAD54B* double KO cells expressing eGFP-RAD51AP1 (Supplemental Fig. S4F, lane 8). We then prepared protein lysates from *RAD54*/*RAD51AP1* double KO cells transiently expressing eGFP-RAD51AP1 under unperturbed conditions (NT), and at 4 and 20 h after release from a 2-h treatment with 0.5 µM MMC. RAD54B was present in anti-eGFP complexes from both untreated and MMC-treated cells (Fig. 4E, lanes 6-7; Supplemental Fig. S4G, lanes 7-8). These results show that endogenous RAD54B can associate with ectopically expressed RAD51AP1 in *RAD54/RAD51AP1* double KO cells in the absence and in the presence of MMC-induced DNA damage.

### Deletion of both RAD51AP1 and RAD54 synergistically sensitizes HeLa cells to olaparib

HR deficiency selectively confers sensitivity to PARPi (Bryant et al. 2005; Farmer et al. 2005). Hence, we asked if single and *RAD54*/*RAD51AP1* double KO cells were sensitive to treatment with the PARPi olaparib. Compared to HeLa cells, *RAD51AP1* KO cells showed mildly increased sensitivity to olaparib (*p* < 0.001), while both *RAD54* KO cell lines (KO-1 and KO-2) were more sensitive (*p* < 0.05 and *p* < 0.01 for KO-1 and KO-2 compared to *RAD51AP1* KO cells, respectively; Fig. 5A). Combined loss of both *RAD51AP1* and *RAD54* further sensitized cells to olaparib (*p* < 0.05 for *RAD51AP1*/*RAD54* KO-1 and KO-2 cells compared to the *RAD54* KO; Fig. 5A). Next, we compared the cytotoxicity to olaparib of *RAD54/RAD51AP1* double KO cells to that of the *RAD54/RAD54B* double KOs. Treatment with olaparib decreased the survival of both *RAD54*/*RAD51AP1* and *RAD54*/*RAD54B* double KO cells to similar extent (*p* < 0.0001 compared to HeLa cells; Fig. 5B).

**Figure 5.**
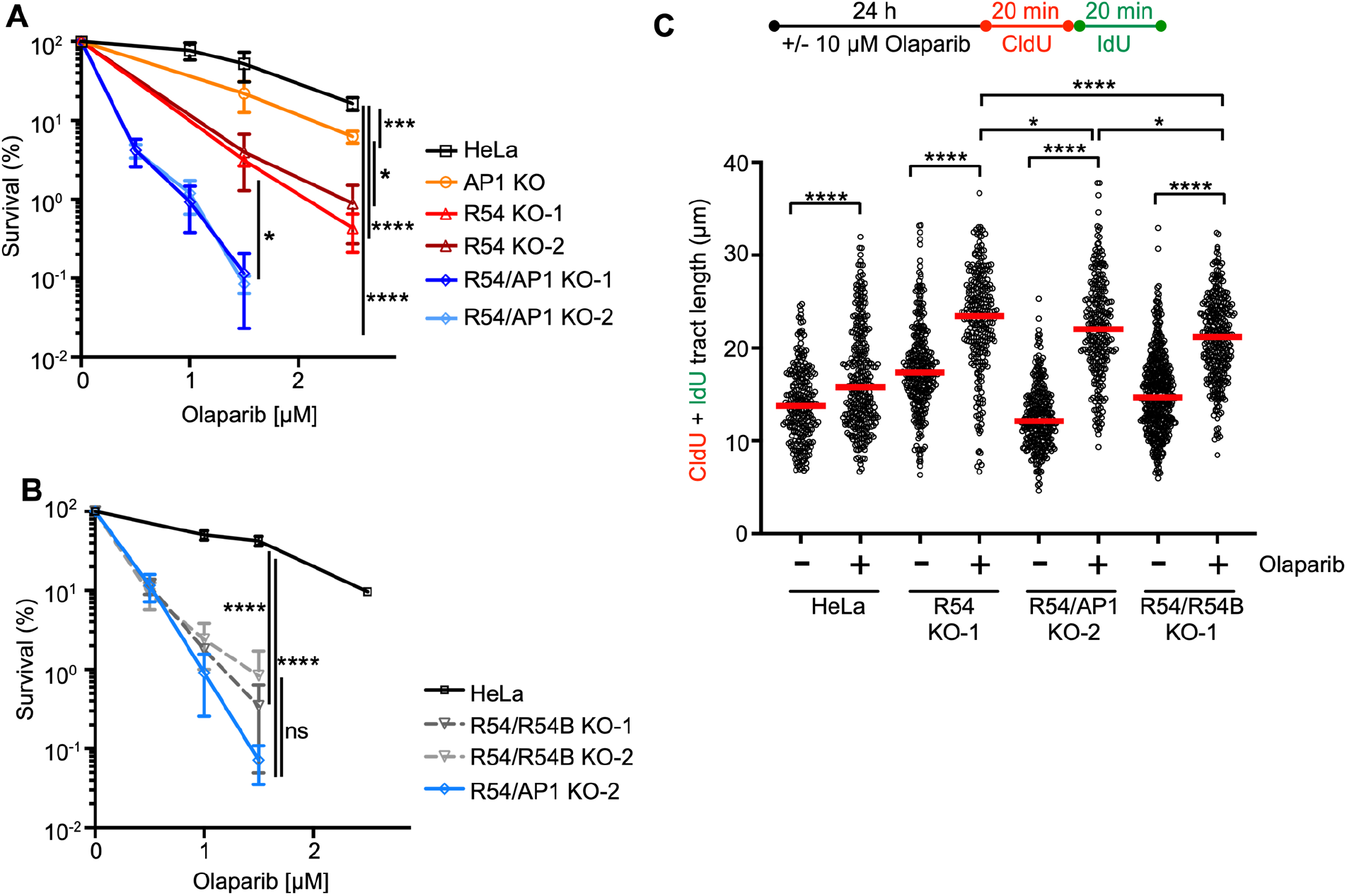
Loss of *RAD54* and *RAD51AP1* or of *RAD54* and *RAD54B* synergistically enhances cellular sensitivity to olaparib. (A) Results from clonogenic cell survival assays in the presence of olaparib of HeLa, *RAD51AP1* KO (AP1 KO), *RAD54* KO (R54 KO-1, KO-2), and *RAD54*/*RAD51AP1* double KO (R54/AP1 KO-1, KO-2) cells. Data points are the means from two independent experiments ±SD. *, *p* < 0.05; ***, *p* < 0.001; ****, *p* < 0.0001; ns, non-significant; two-way ANOVA test followed by Tukey’s multiple comparisons test. (B) Results from clonogenic cell survival assays in the presence of olaparib of HeLa, *RAD54*/*RAD54B* double KO (R54/R54B KO-1, KO-2), and *RAD54*/*RAD51AP1* double KO (R54/AP1 KO-2) cells. Data points are the means from two independent experiments ±SD. ****, *p* < 0.0001; ns, non-significant; two-way ANOVA test followed by Tukey’s multiple comparisons test. (C) Schematic of the experimental protocol for the DNA fiber assay in unperturbed cells and after olaparib. Total tract length (CldU+IdU) of DNA fibers from HeLa, *RAD54* KO (R54 KO-1), *RAD54*/*RAD51AP1* double KO (R54/AP1 KO-2), and *RAD54*/*RAD54B* double KO (R54/R54B KO-1) cells with and without olaparib. Data points are from 130-230 fibers of two independent experiments each, with medians (red lines). *, *p* < 0.05; ****, *p* < 0.0001; Mann-Whitney test.

To understand why, in comparison to *RAD54*/*RAD54B* double KO cells, *RAD54*/*RAD51AP1* double KO cells respond with significantly increased sensitivity to MMC and HU albeit with similar sensitivity to olaparib, we analyzed the dynamics of replication fork progression by DNA fiber assay after a 1-day incubation of cells in 10 µM olaparib. Compared to untreated cells, fiber tracts were longer in HeLa cells after olaparib (Fig. 5C; Supplemental Fig. S5A; Table S1), consistent with the results from an earlier study (Maya-Mendoza et al. 2018). In *RAD54* single, and *RAD51AP1*/*RAD54* and *RAD54/RAD54B* double KO cells, fiber tracts were significantly longer than in HeLa cells (*p* < 0.0001; Fig. 5D; Supplemental Fig. S5A; Table S1), indicative of the further increased defects of the KO cell lines in restraining fork progression after olaparib. Compared to the lengths of fiber tracts obtained under unperturbed conditions, median fiber tracts were 15% longer in HeLa cells, 35% longer in *RAD54* KO cells and 82% and 44% longer in *RAD54/RAD51AP1* and *RAD54/RAD54B* double KO cells, respectively. Collectively, these results show that HR-proficient HeLa cells restrain accelerated fork elongation more effectively than *RAD54* single and *RAD54*/*RAD51AP1* and *RAD54*/*RAD54B* double KO cells. Moreover, while a 1-day exposure to olaparib is associated with increased levels of DSBs in all cell lines investigated, COMET assays revealed significantly more olaparib-induced DSBs in *RAD54*/*RAD51AP1* and *RAD54*/*RAD54B* double KO cells than in HeLa cells and the *RAD54* single KO (*p* < 0.0001; Supplemental Fig. SC; Table S2). These results suggest that fork stability is particularly compromised when fork movement is accelerated in *RAD51AP1*/*RAD54* and *RAD54*/*RAD54B* double KO cells, and that after olaparib the stress to replication forks, as determined by COMET assay, is similar in *RAD51AP1*/*RAD54* and *RAD54*/*RAD54B* double KO cells and higher than in HeLa cells and the *RAD54* single KO.

## Discussion

In this study, we have shown that the HR function of RAD54 can largely be compensated for by the RAD51AP1 protein. Surprisingly, in the context of stalled and collapsed DNA replication (after HU or MMC), the compensatory activity of RAD51AP1 is greater than that of RAD54B (Fig. 6A). After treatment of cells with olaparib, however, RAD51AP1 and RAD54B are equally important in substituting for RAD54, which may stem from the similar degree of engagement of both proteins in post-replication repair (Fig. 6B).

**Figure 6.**
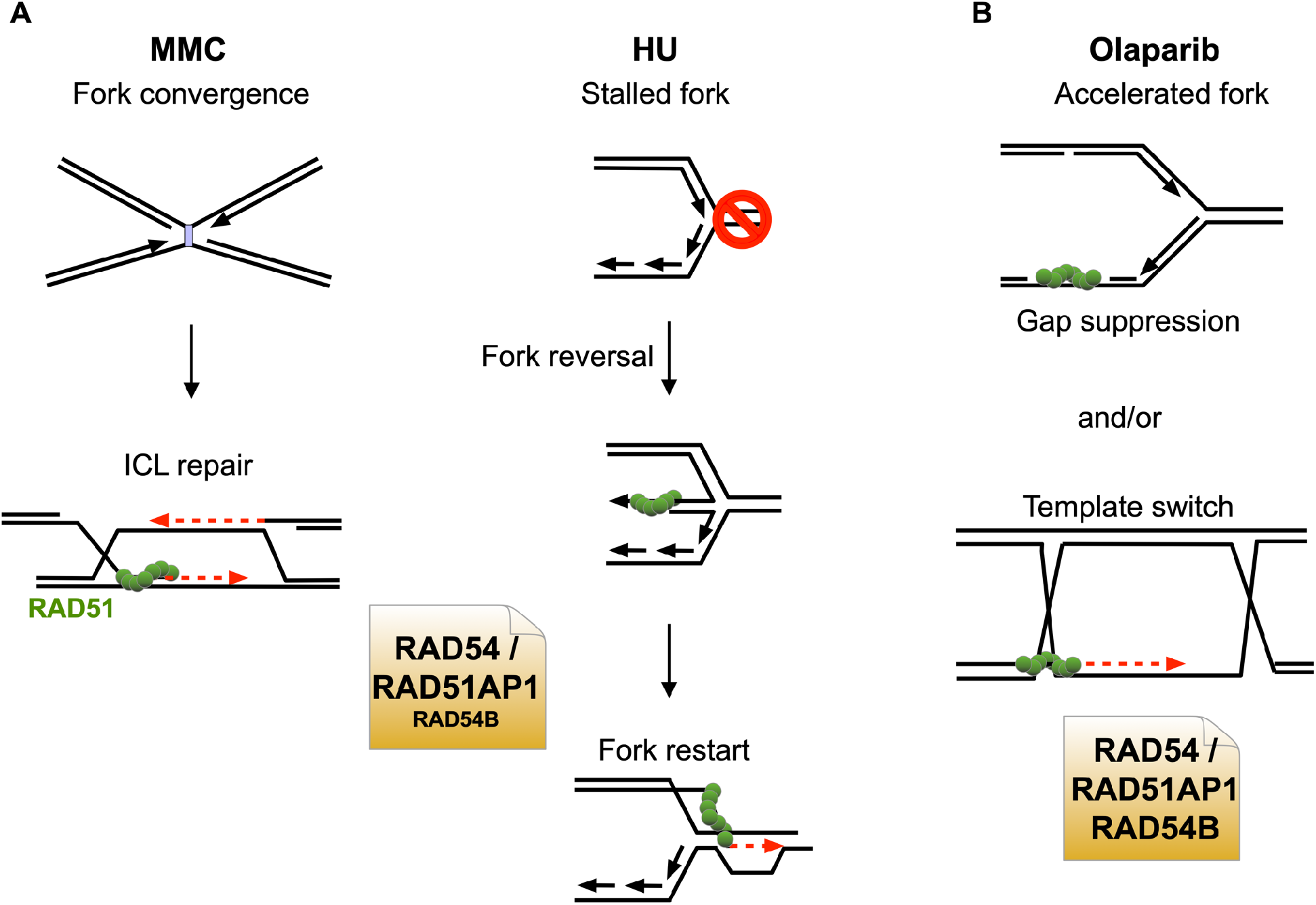
Model depicting the predominant synthetic interactions between RAD54 and RAD51AP1 (A, B) and between RAD54 and RAD54B (B). For details see text.

Given that a complex between RAD51AP1 and RAD54B can be isolated (this study), evidence of physical interaction and functional cooperation between purified RAD54B and RAD51 (Sarai et al. 2006; Wesoly et al. 2006), and of an indirect association between human RAD54B and RAD51 in the context of chromatin and in cells (Tanaka et al. 2000; Zhang et al. 2007), it is possible that in select stages of the HR reaction, within a certain context of the genome, or in response to specific types of DNA damage, RAD51AP1 may function cooperatively with RAD54B, possibly bridging RAD54B to RAD51. Of note, *RAD54* KO cells with RAD54B knockdown show increased sensitivity to MMC while RAD54B-depleted *RAD54*/*RAD51AP1* double KO cells do not. These results could suggest that RAD51AP1 may be required for RAD54B function. However, *RAD54*/*RAD51AP1* double KO cells are significantly more sensitive to MMC than *RAD54*/*RAD54B* double KO cells, which argues that RAD51AP1 has additional function(s) aside from working with RAD54B in repairing MMC-induced DNA damage. This could be together with another and yet to be identified translocase, or within transcriptionally active, decondensed regions of the genome in which RAD51AP1 specifically was shown to promote HR (Ouyang et al. 2021).

PARPi treatment leads to the formation of replication associated ssDNA gaps (Maya-Mendoza et al. 2018; Thakar and Moldovan 2021). Gap suppression mechanisms and HR-mediated post-replicative repair serve to restrict and eliminate ssDNA replication gaps (Hashimoto et al. 2010; Piberger et al. 2020; Cong et al. 2021; Panzarino et al. 2021). We speculate that concomitant loss of RAD54 and either RAD51AP1 or RAD54B may exacerbate ssDNA gap formation, inhibiting the annealing of nascent DNA strands and, as such, the reversal of replication forks (Cong et al. 2021), resulting in the similar degree of synthetic lethality between *RAD54* and *RAD51AP1* or *RAD54* and *RAD54B* with PARPi (Fig. 6B).

RAD51AP1 expression is increased in different breast cancer subtypes and other cancers and inversely associated with overall survival (Song et al. 2004; Henson et al. 2006; Martin et al. 2007; Martinez et al. 2007; Obama et al. 2008; Pathania et al. 2016; Chudasama et al. 2018; Li et al. 2018; Bridges et al. 2020; Zhao et al. 2020; Zhuang et al. 2020). Moreover, *Rad51ap1* deficiency abrogates tumor growth and metastasis in a breast cancer mouse model (Bridges et al. 2020), suggesting that the RAD51AP1 protein may be a promising target of inhibition in anti-cancer therapy. Given our results showing extensive redundancy between RAD51AP1 and RAD54, we surmise that the simultaneous inactivation of both RAD51AP1 and RAD54 would be a viable chemotherapeutic strategy to treat cancer. Targeting RAD51AP1 together with RAD54 may be particularly effective against tumors with overactive HR (Raderschall et al. 2002; Xu et al. 2005; Klein 2008; Marsden et al. 2016), cancerous cells maintaining their telomeres by the ALT pathway (Barroso-Gonzalez et al. 2019; Mason-Osann et al. 2020; Recagni et al. 2020), and BRCA1/2-mutant tumors that have regained HR proficiency and are resistant to PARPi (Han et al. 2020; Kim et al. 2021).

## Materials and methods

### Cell culture, transfections, and siRNAs

HeLa and A549 cells were obtained from ATCC and were maintained as recommended. HeLa cells in which either *RAD51AP1* or *RAD54* are deleted were maintained as described previously (Liang et al. 2019; Maranon et al. 2020). SiRNAs were obtained from Qiagen (Supplemental Table S3). SiRNA forward transfections with Lipofectamine RNAiMAX (Invitrogen) were performed on two consecutive days. The concentration of siRNAs in transfections was 20 nM each. Cells were treated with drugs at 96 h after the first transfection.

### Generation of RAD54/RAD51AP1 and RAD54/RAD54B double KO cells

*RAD51AP1* knockout (KO) and *RAD54* KO HeLa cells, that we described previously (Liang et al. 2019; Maranon et al. 2020), were used to generate *RAD54/RAD51AP1* and *RAD54*/*RAD54B* double KO cells. Briefly, a combination of two *RAD54* or *RAD54B* CRISPR/Cas9n(D10A) KO plasmids each containing one of two different sgRNAs (*i*.*e*., sgRNA (54)-A and sgRNA (54)-B; sgRNA (54B)-A and sgRNA (54B)-B; Supplemental Table S3) was purchased from Santa Cruz Biotechnology (sc-401750 for *RAD54*; sc-403794 for *RAD54B*) and used to transfect single KO cells as described (Maranon et al. 2020). Disruption of *RAD54* and *RAD54B* was validated by sequence analysis after genomic DNA was isolated from a selection of edited and non-edited clonal isolates using DNeasy Blood & Tissue Kit (Qiagen). *RAD54* and *RAD54B* genomic DNA sequences were amplified by PCR using primer pairs flanking the sgRNA target sites (Supplemental Table S3). PCR products were gel purified, cloned into pCR4-TOPO (Invitrogen) and transformed into TOP10 competent *E. coli*. Plasmid DNA was prepared using ZR Plasmid Miniprep-Classic Kit (Zymo Research) and submitted for Sanger sequencing.

### Generation of RAD54 expressing RAD51AP1 KO, RAD54 KO and RAD54/RAD51AP1 DKO cells

The plasmid containing the C-terminally HA-tagged full-length human RAD54 cDNA has been described (Maranon et al. 2020). A *Kpn*I to *Not*I digest was performed to clone RAD54-HA into pENTR1A (Invitrogen), followed by transfer into pLentiCMV/TO DEST#2 (Campeau et al. 2009) using Gateway LR Clonase II (Invitrogen) for the production of lentiviral particles in HEK293FT cells (Invitrogen), as described (Campeau et al. 2009). Lentivirus was used to transduce *RAD51AP1* KO, *RAD54* KO, and *RAD54*/*RAD51AP1* double KO cells in 6 µg/ml polybrene, as described (Campeau et al. 2009).

### Clonogenic cell survival assays and Western blot analysis

Clonogenic cell survival assays after mitomycin C (MMC; Sigma) were performed as described (Maranon et al. 2020). To assess cellular sensitivity to olaparib (AZD2281; KU-0059436; Selleck Chemicals), cells were chronically exposed to 0.5-2.5 μM olaparib in regular growth medium for 12 days, as described (Spies et al. 2016). Cells were fixed and stained with crystal violet to determine the fraction of cells surviving.

Western blot analyses were performed according to our standard protocols (Wiese et al. 2006). The following primary antibodies were used: α-RAD51AP1 ((Dray et al. 2010); 1:6,000), α-RAD54 (F-11; sc-374598; Santa Cruz Biotechnology; 1:1,000); α-RAD51 (Ab-1; EMD Millipore; 1:3,000), α-PARP1 (ab6079; Abcam; 1:2,000), α-β-Actin (ab8226;Abcam; 1:1,000), α-HA.11 (MMS-101R; BioLegend; 1:1,000) and α-RAD54B ((Wesoly et al. 2006); 1:1,000). HRP-conjugated goat anti-rabbit or goat anti-mouse IgG (Jackson ImmunoResearch; 1:10,000) were used as secondaries. Western blot signals were acquired using a Chemidoc XRS+ gel imaging system and Image Lab software version 5.2.1 (BioRad).

### Cell cycle analysis and flow cytometry

Cell cycle analysis and flow cytometry were performed as described (Maranon et al. 2020), except that exponentially growing cells were treated with 0.5 μM MMC for 2 hours, washed twice with warm PBS and incubated in fresh growth medium for the times indicated prior to pulse-labeling with 10 μM EdU.

### Metaphase spreads

For the assessment of chromosomal aberrations, 2×10^5^ cells were seeded in 6-well tissue culture plates and incubated at 37°C for 24 hours before exposure to 4 mM hydroxyurea (HU; Sigma) in regular growth medium for 5 hours, as described (Schlacher et al. 2011). After HU treatment, cells were washed in warm PBS and incubated in medium containing 0.1 μg/ml colcemid (SERVA) for 24 hours. Cells were detached and allowed to swell in 0.075 M KCl at 37°C for 30 minutes and fixed in methanol:acetic acid (3:1), as described (Parplys et al. 2015). Cells were dropped onto wet slides, air dried and stained in 3% Giemsa in Sorensen buffer (0.2 M Na2HPO4/NaH2PO4, pH7.3) at room temperature for 10 min. Images were acquired using Zeiss Axio-Imager.Z2 microscope equipped with Zen Blue software (Carl Zeiss Microscopy) using a 63x oil objective. One hundred metaphases were assessed per sample.

### DNA fiber assay

DNA replication progression was assessed by the single-molecule DNA fiber assay and essentially as described previously (Schlacher et al. 2011; Parplys et al. 2015; Taglialatela et al. 2017). Briefly, exponentially growing cells were pulse-labelled in regular growth medium containing 25 μM CldU for 20 min, followed by a 5-h incubation in regular growth medium with 4 mM HU, after which the cells were pulse-labelled in regular growth medium containing 250 μM IdU for 20 min. Cells were detached from the cell culture dish by scraping in ice-cold PBS, adjusted to 4×10^5^ cell/ml and processed for fiber spreading as described (Parplys et al. 2015). In a modified version of this assay, cells were exposed for 24 h in 10 μM olaparib, followed by two consecutive rounds of 20 min each in CldU first and then in IdU (Maya-Mendoza et al. 2018). Images were acquired using Zeiss Axio-Imager.Z2 microscope equipped with Zen Blue software (Carl Zeiss Microscopy) using a 63x oil objective. Per sample and condition 200 fiber tracts were measured using ImageJ software (https://imagej.net).

### Co-immunoprecipitations

The peGFP-RAD51AP1 expression vector is based on peGFP-C1 (Clontech) and has been described previously (Modesti et al. 2007). *RAD54*/*RAD51AP1* double KO cells were transfected with peGFP-C1 or peGFP-RAD51AP1 and Lipofectamine2000 (Invitrogen). Twenty-four hours after transfection, cells were subjected to a medium change or treated with 0.5 µM MMC for 2 hours. Cells were washed twice with warm PBS, fresh medium was added, and cells were incubated for the times indicated. Cells were lysed in chilled lysis buffer containing 50 mM Tris-HCl, pH 7.5, 300 mM NaCl, and 0.5% NP-40, supplemented with EDTA-free protease inhibitor cocktail (Roche) and HALT phosphatase inhibitors (Thermo Fisher Scientific). For 1.5×10^6^ cells, 25 μl of GFP-Trap ® dynabeads (ChromoTek) were used to trap the ectopic proteins. Protein lysates were diluted to 50 mM Tris-HCl, pH 7.5, 150 mM NaCl, 0.1% NP-40, and 0.1unit DNase I (Gold Biotechnology) per µg protein, and mixed with the equilibrated beads at 4°C for 1 h with gentle rotation. The GFP-Trap ® dynabeads were washed three times with 500 µl binding buffer, bound protein complexes were eluted in 40 µl 2× LDS buffer (Thermo Fisher Scientific) and fractionated on 7% NuPAGE Tris-Acetate gels (Thermo Fisher Scientific) and for Western blot analysis.

### Statistics and reproducibility

GraphPad Prism 9 software was used to perform statistical analyses. Data obtained are from 2-4 independent experiments, as indicated. To assess statistical significance two-way or one-way ANOVA tests were performed. *P* ≤ 0.05 was considered significant.

## Data availability

All data supporting the findings of this study are available here and in the Supplemental Material.

## Competing interest statement

The authors declare no competing interests.

## Acknowledgements

The authors wish to thank the CSU Flow Cytometry and Cell Sorting Facility for their help in method optimization and sample analyses. This work was supported by a CSU CVMBS College Research Grant and by National Institutes of Health Grants R01 ES021454, R56 ES021454, R03 ES029206 (to C.W.) and R01 ES007061, R35 CA241801 (to P.Su.).

## Author contributions

Y.K., P.Su. and C.W. conceptualization; P.Se., N.S., Y.K. and M.U. data curation; P.Se., N.S., Y.K. and M.U. data analyses and validation; P.Se., N.S., Y.K., P.Su. and C.W. writing of the manuscript; P.Su. and C.W. funding acquisition.

## Notes

### Competing Interest Statement

The authors have declared no competing interest.

### Summary of Updates

Add replicate data in Fig.5 and Fig. S5; editorial changes in the text, change of title.

