## Supplementary Information for "RAD51AP1 and RAD54 underpin two distinct RAD51-dependent routes of DNA damage repair via homologous recombination"

**Supplemental Materials and Methods**

**Supplemental Figure S1** – related to Figs. 1&2

**Supplemental Figure S2** – related to Fig. 2

**Supplemental Figure S3** – related to Fig. 3

**Supplemental Figure S4** – related to Fig. 4

**Supplemental Figure S5** – related to Fig. 5

**Supplemental Table S1**

**Supplemental Table S2**

**Supplemental Table S3**

#### Supplemental Materials and Methods

*Cell fractionation and cell growth assay*

Cell fractionation was carried out using the Subcellular Protein Fractionation Kit (Thermo Fisher Scientific, Waltham, MA) and as described by the manufacturer. To determine the rates of cell growth, cells were seeded in 6-well tissue culture plates at a density of $1.2x10^4$ cells/well. Cell were counted after trypsinization for eight consecutive days. Cell numbers were plotted using nonlinear regression, and population doubling times (PDL) were calculated using the following equation:

$$\mathrm{PDL} = 3.32 \times \log\left(\frac{\text{total viable cells at harvest}}{\text{total viable cells at seed}}\right)$$

*Recombination assay*

U2OS-DRGFP (*i.e.* DR-U2OS) cells have been described elsewhere (Nakanishi et al. 2005; Xia et al. 2006). The gene conversion assay using U2OS-DRGFP cells was performed as previously described (Parplys et al. 2015; Liang et al. 2016; Liang et al. 2020). Briefly, $1.5\times10^5$ cells were seeded in 6-well tissue culture plates 24 hours prior to transfection in antibiotic free media. I-*Sce*I was expressed transiently in U2OS-DRGFP cells from the pCβASCE expression vector (Richardson et al. 1998) at 0.8 µg/150,000 cells and co-transfected with siRNA using Lipofectamine2000 (Invitrogen). Transfected cells were kept in regular growth medium for 72 hours, after which time they were analyzed by flow cytometry to measure the percentage of viable cells expressing GFP, as previously described (Parplys et al. 2015). To assess the fraction of GFP-positive cells, FlowJo version 10.7 (BD Biosciences) was used.

*Indirect immunostaining, microscopy, and image analysis*

Indirect immunostaining was performed as described (Maranon et al. 2020). The primary antibody used to detect RAD54 foci was α-RAD54 (F-11; sc-374598; Santa Cruz Biotechnology; 1:1,000). A Zeiss Axio-Imager.Z2 microscope equipped with Zen Blue software (Carl Zeiss Microscopy) was used to take images using a 63x oil objective. In each channel, 18 Z-stacks were obtained as 0.2 µm slices. Images were processed in Fiji (https://imagej.net/Fiji), separating the channels and producing maximum projection files.

*Immunoprecipitations*

*RAD51AP1* KO cells stably expressing FLAG-tagged RAD51AP1 were used. *RAD51AP1* was introduced into *RAD51AP1* KO cells by transduction with lentivirus, as described previously (Campeau et al. 2009). Protein lysate from HeLa with endogenous RAD51AP1 (control) and from *RAD51AP1* KO cells expressing FLAG-RAD51AP1 were prepared in chilled buffer containing 50 mM Tris-HCl, pH 7.5, 300 mM NaCl, and 0.5% NP-40 supplemented with EDTA-free protease inhibitor cocktail (Roche) and HALT phosphatase inhibitors (ThermoScientific). Protein lysates were adjusted to 150 mM NaCl, 0.1% NP-40, and 0.1 unit/µg protein DNase I (GoldBio). Twenty-five µl ANTI-FLAG ® M2- resin (Sigma) was first equilibrated with binding buffer (50 mM Tris-HCl, pH 7.5, 150 mM NaCl and 0.1% NP-40) before protein lysates containing 2 mg total protein were added and incubated for at 4°C for 1 hour with gentle rotation. Protein complexes bound to anti-FLAG resin were washed three times with 500 µl binding buffer, and bound protein complexes were eluted with 150 ng/µl 3× FLAG peptide (Sigma) in binding buffer. Eluted protein was fractionated on 7.5% NuPAGE Tris-Acetate gels, transferred onto polyvinylidene fluoride membrane (Millipore), and detected by Western blot analysis.

*Comet assay*

The Comet assay (Trevigen) was completed according to the manufacturer's instructions. Briefly, $5\times10^4$ cells were seeded in 60-mm plates. Cells were incubated at 37°C for 48 h before treated with 10 µM olaparib or DMSO for 24 h. Cells were harvested in cold PBS, counted and combined with molten low melting (LM)agarose at 1:10 to $1\text{x}10^5$ cells/ml. Fifty µl of the suspension were transferred on to COMET slides and incubated in the dark at 4°C for 30 minutes. Slides were incubated in cold lysis buffer at 4°C overnight. Excess lysis buffer was drained and electrophoresis was performed in cold neutral COMET electrophoresis buffer (10 mM Tris base, 250 mM sodium acetate) at 25 V and 4°C for 30 min. The slides were immersed in DNA precipitation solution (Trevigen) for 30 min, followed by incubation in 70% ethanol for 30 min. Slides were the dried at 37°C for 30 min and stained with 0.3× SYBR Gold (Thermo Fisher). Images were acquired on a Zeiss Axio-Imager.Z2 microscope equipped with Zen Blue software (Carl Zeiss Microscopy) using a 20× objective, and 100 comets were measured per condition. The length comet tails were measured using ImageJ software (https://imagej.net).

### Supplemental Figures

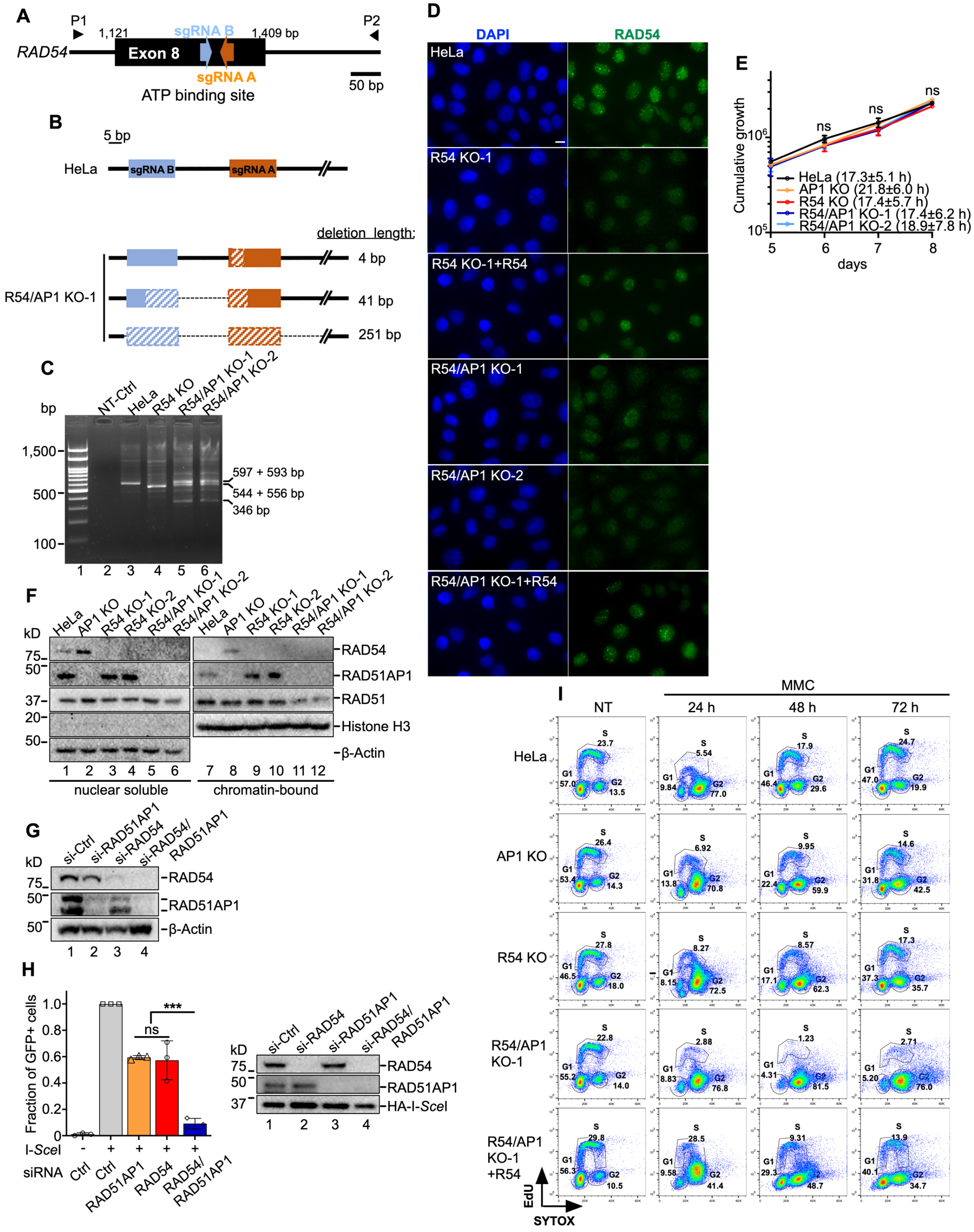

**Supplemental Figure S1**. (*A*) Schematic of exon 8 and adjacent introns of the human *RAD54* gene with the location of the sgRNA targeting sequences indicated in orange and blue (for sequences of sgRNAs see Table S3) and of PCR primer pair (P1, P2) indicated as arrow heads (for sequences of PCR primers see Table S3). (*B*) Schematics of the altered sequences of obtained PCR amplicons and deletions in *RAD54* detected in *RAD54*/*RAD51AP1* KO-1 cells after sequence analyses. (*C*) Representative agarose gel of PCR products obtained from genomic DNA of HeLa, *RAD54* KO and the two *RAD54*/*RAD51AP1* KO cell lines using primers P1 and P2. NT-Ctrl: non-template control. (*D*) Representative micrographs of RAD54 foci (green) in HeLa, *RAD54* KO-1, *RAD54*/*RAD51AP1* double KO (KO-1 and KO-2) cells, and in *RAD54* KO-1 and *RAD54*/*RAD51AP1* KO-1 cells stably expressing RAD54-HA (here: +R54); scale bar = 10 µm. (*E*) Growth curves under unperturbed conditions of HeLa and KO cell lines. Data points are the means from three experiments ±SD. ns, non-significant; two-way ANOVA test followed by Tukey's multiple comparisons test. (*F*) Western blots of soluble and chromatin-bound nuclear extracts of HeLa cells and single and double KO cell lines. The signals for β-Actin and histone H3 serve as a loading and fractionation control, respectively. (*G*) Western blots of whole cell protein lysates to show the extent of RAD54/RAD51AP1 protein knockdown in A549 cells. Loading control: β-Actin. (*H*) Left: Average percentage of GFP-positive cells normalized to non-targeting control siRNA (Ctrl) after RAD51AP1 and/or RAD54 knockdown in U2OS-DRGFP cells. Right: Western blots of nuclear extracts to show the extent of RAD54/RAD51AP1 protein knockdown and I-*Sce*I expression in U2OS-DRGFP cells. Bars are the means from three independent experiments ±SD. Symbols are data points from individual experiments. ***, $p<0.001$; ns, non-significant; one-way ANOVA test followed by Tukey's multiple comparisons test. (*I*) Representative results from flow cytometry showing two-color fluorescence of cell cycle profiles from the cells used in Fig. 1D and Fig. 2B. Y-axis: EdU, X-axis: SYTOX.

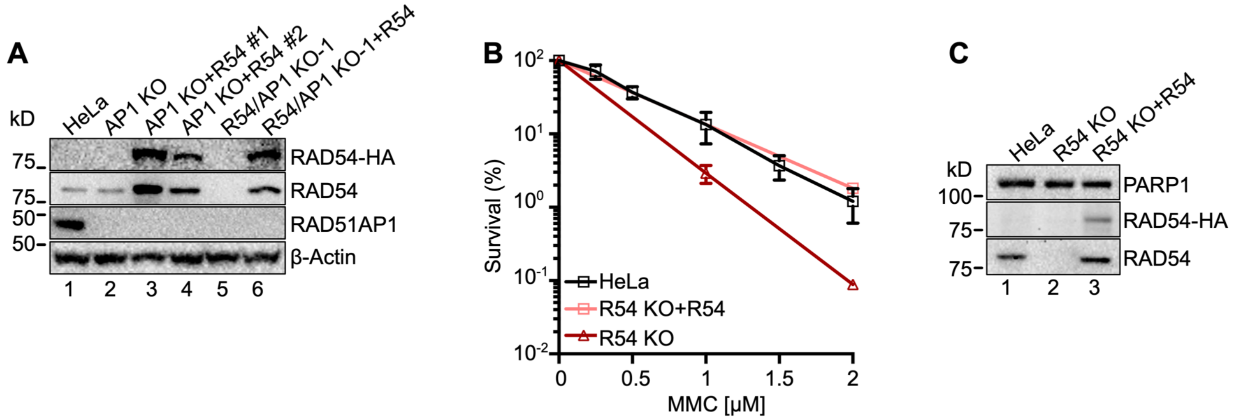


**Supplemental Figure S2**. (*A*) Western blots of whole cell protein extracts to show stably expressed ectopic RAD54-HA (here: +R54) in *RAD51AP1* KO (AP1 KO; lanes 3 and 4) and *RAD54*/*RAD51AP1* double KO (R54/AP1 KO-1; lane 6) cells. Loading control: β-Actin. (*B*) Results from MMC clonogenic cell survival assays of HeLa cells and *RAD54* KO (R54 KO) cells with and without ectopic RAD54-HA (here: +R54). Data points for *RAD54* KO+RAD54-HA cells are the means from three technical replicates. Data points for *RAD54* KO and HeLa cells are the means from three independent experiments ±SD. (*C*) Western blots of whole cell protein extracts to show stably expressed ectopic RAD54-HA in *RAD54* KO cells. Loading control: PARP1.

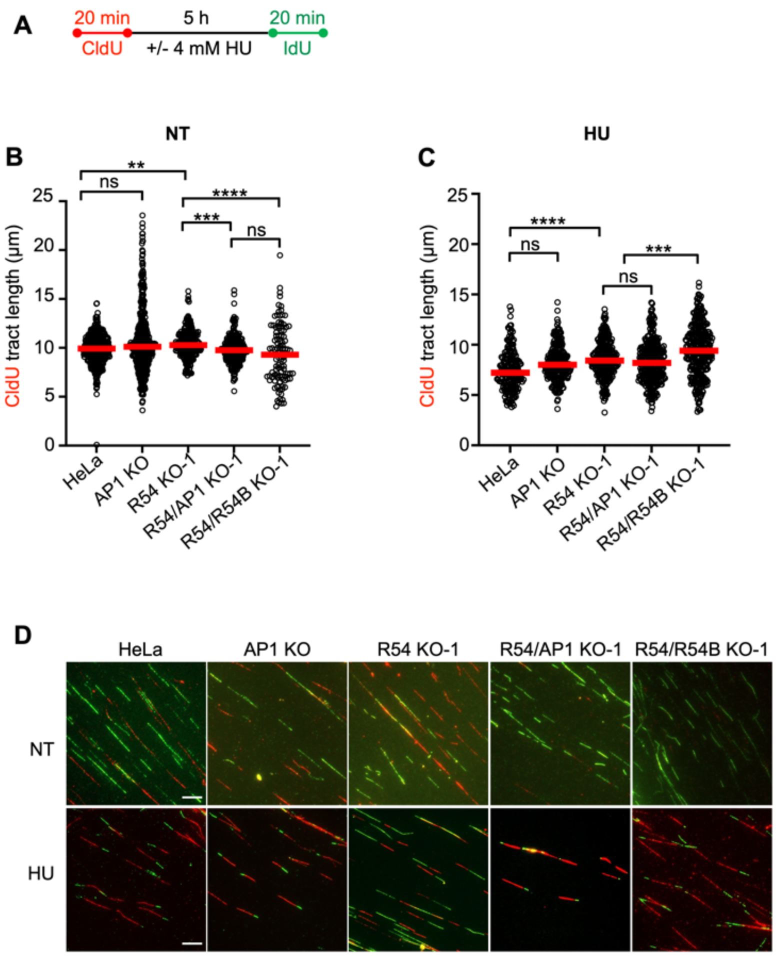


**Supplemental Figure S3**. (*A*) Schematic of experimental protocol for the DNA fiber assay with and without HU and the data presented in B-D. (*B, C*) Median CldU (red) tract lengths in cells without (NT) or with HU treatment. Data points are from 100-150 fibers of two independent experiments each, with medians (red lines). ****, $p$<0.0001; ***, $p$<0.001; **, $p$<0.01; ns, non-significant; Kruskal-Wallis test followed by Dunn's multiple comparisons test. (*D*) Representative micrographs of DNA fibers from HeLa, single, and double KO cells; scale bars = 10 µm.

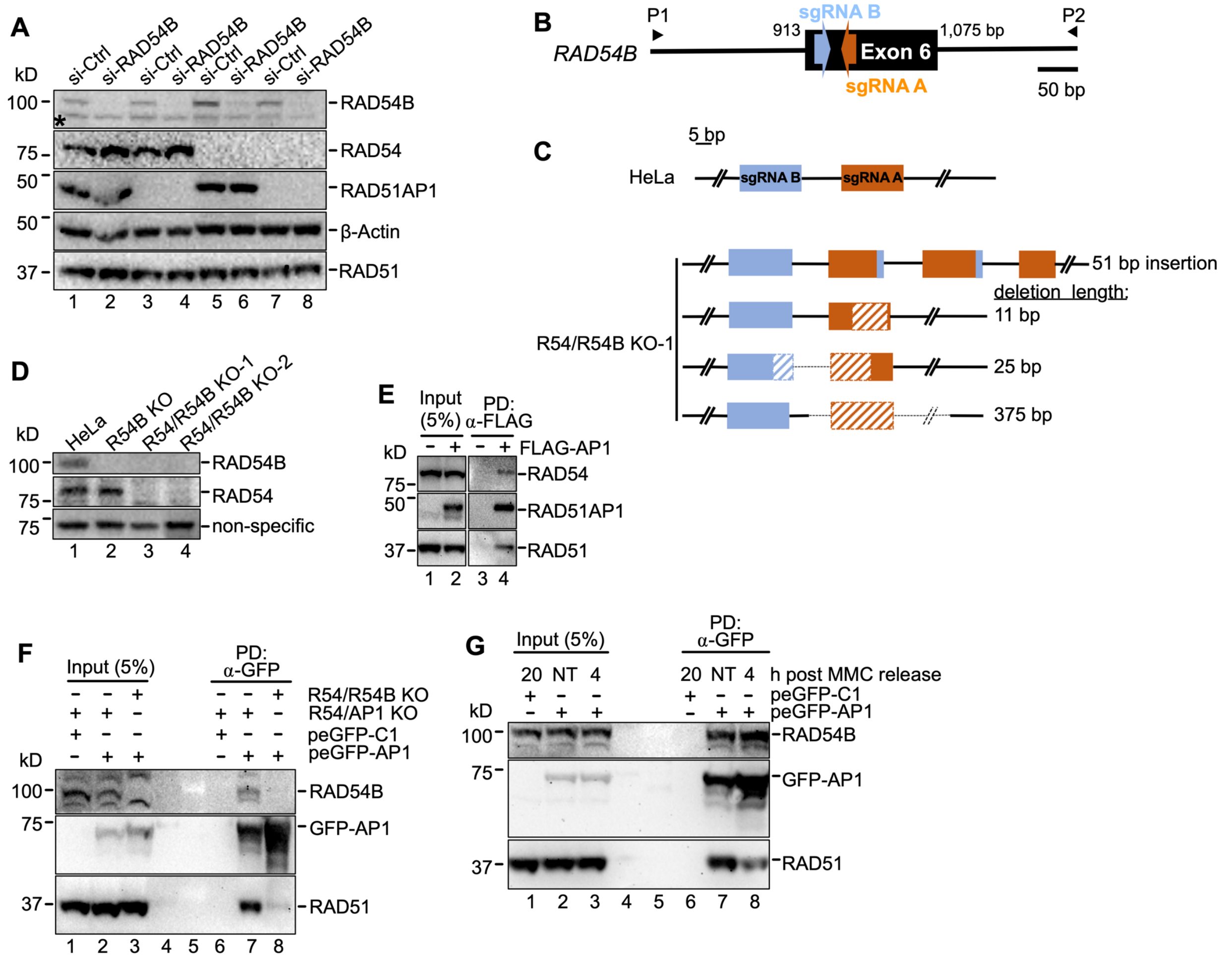


**Supplemental Figure S4**. (*A*) Western blots of whole cell protein extracts of HeLa, *RAD51AP1* KO (AP1 KO), *RAD54* KO (R54 KO-1) and *RAD54/RAD51AP1* (R54/AP1 KO-1) double KO cells. Loading control: β-Actin. (*B*) Schematic of exon 6 and adjacent introns of the human *RAD54B* gene with the location of the sgRNA targeting sequences (in orange and blue; see Table S3) and PCR primers P1 and P2 (see Table S3) used for amplification. (*C*) Schematics of the altered sequences (including one insertion and three deletions) of obtained PCR products from four *RAD54B* alleles detected in *RAD54/RAD54B* KO-1 cells after sequence analyses. (*D*) Western blots of whole cell protein extracts to show loss of expression of both RAD54B and RAD54 in two independently isolated *RAD54/RAD54B* double KO cell lines (KO-1 and KO-2; lanes 3 and 4). Loading control: non-specific band detected after probing with RAD54B antibody. (*E*) Western blots to show that endogenous RAD54 co-precipitates in anti-FLAG pull-downs (PD) of ectopically expressed FLAG-RAD51AP1 in *RAD51AP1* KO cells (lane 4). The signal for RAD51 serves as a positive control. (*F*) Western blots to show that endogenous RAD54B co-precipitates in anti-GFP pull-downs (PD) of ectopically expressed GFP-RAD51AP1 in *RAD54*/*RAD51AP1* double KO cells (lane 7), but not in anti-GFP PDs of ectopically expressed GFP-RAD51AP1 in *RAD54*/*RAD54B* double KO cells (lane 8). The signal for RAD51 serves as a positive control. Note: Diminished RAD51 is detected in anti-GFP PDs of GFP-RAD51AP1 expressed in *RAD54*/*RAD54B* double KO cells (lane 8), possibly due to endogenous RAD51AP1 interfering with complex formation between GFP-RAD51AP1 and RAD54B. (*G*) Western blots to show that endogenous RAD54B co-precipitates in anti-GFP pull-downs (PD) of ectopically expressed GFP-RAD51AP1 in *RAD54*/*RAD51AP1* double KO cells, constitutively and at 4 h post release from a 2-h incubation of cells in 0.5 μM MMC (lanes 7 and 8, respectively).

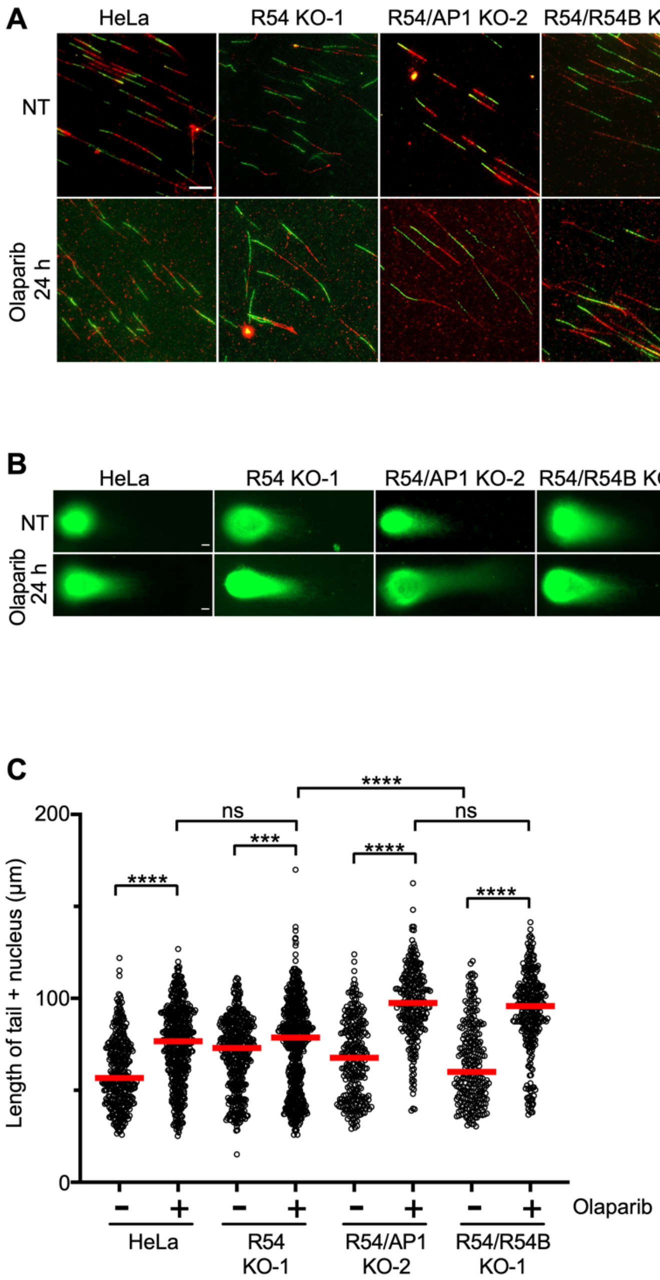


**Supplemental Figure S5**. (*A*) Representative micrographs of DNA fibers in unperturbed cells (NT) and after a 24-h incubation in 10 µM olaparib. Scale bar = 10 µm. (*B*) Representative micrographs of nuclei with eluted DNA after COMET assay of unperturbed cells (NT) and after a 24-h incubation in 10 µM olaparib. Scale bars = 10 µm. (*C*) Quantification of COMET assay of unperturbed cells (NT) and after a 24-h incubation in 10 µM olaparib. Data points are from 120-230 nuclei of 2-3 independent experiments each. ****, $p < 0.0001$; ***, $p<0.001$; ns, non-significant; Mann-Whitney test.

#### Supplemental Tables

**Table S1. Summary of DNA fiber data analyses.** Means, medians, 25th and 75th percentiles, *p*-values and number of experiments (N), as shown in the graphs of the corresponding figures.

| Fig. | Label | Genotype | Treat-ment | Mean (µm) | Median (µm) | 25th percentile | 75th percentile | *p* | N |
|---|---|---|---|---|---|---|---|---|---|
| **3C** | IdU | WT | HU | 6.33 | 6.06 | 5.07 | 7.41 | - | 2 |
| **3C** | IdU | AP1 KO | HU | 5.84 | 5.53 | 4.31 | 6.93 | 0.0758[§] | 2 |
| **3C** | IdU | R54 KO-1 | HU | 7.34 | 6.91 | 5.58 | 8.80 | 0.0036[§] | 2 |
| **3C** | IdU | R54/AP1 KO-1 | HU | 3.26 | 3.16 | 2.29 | 4.11 | <0.0001[§,†] | 2 |
| **3D** | IdU | WT | NT | 10.09 | 9.95 | 9.07 | 10.88 | - | 2 |
| **3D** | IdU | AP1 KO | NT | 10.46 | 10.33 | 9.50 | 11.32 | 0.0044[§] | 2 |
| **3D** | IdU | R54 KO-1 | NT | 9.95 | 10.03 | 8.99 | 11.07 | >0.9999[§] | 2 |
| **3D** | IdU | R54/AP1 KO-1 | NT | 9.59 | 9.42 | 8.53 | 10.53 | <0.0001[§] 0.0002[†] | 2 |
| **4C** | IdU | WT | NT | 10.55 | 10.53 | 9.73 | 11.49 | - | 2 |
| **4C** | IdU | R54/AP1 KO-1 | NT | 9.59 | 9.42 | 8.53 | 10.53 | <0.0001[§] | 2 |
| **4C** | IdU | R54/R54B KO-1 | NT | 10.05 | 10.07 | 9.36 | 10.72 | 0.0104[§] | 2 |
| **4C** | IdU | R54/R54B KO-2 | NT | 10.53 | 10.46 | 9.69 | 11.11 | >0.9999[§] | 2 |
| **4D** | IdU | WT | HU | 5.68 | 5.37 | 4.73 | 6.71 | - | 2 |
| **4D** | IdU | R54/AP1 KO-1 | HU | 3.19 | 3.04 | 2.34 | 4.05 | <0.0001[§,‡] | 2 |
| **4D** | IdU | R54/R54B KO-1 | HU | 5.05 | 4.77 | 3.97 | 5.94 | 0.0007[§] | 2 |
| **4D** | IdU | R54/R54B KO-2 | HU | 4.83 | 4.46 | 3.64 | 5.69 | <0.0001[§] 0.3580[¶] | 2 |
| **5C** | CldU + IdU | WT | NT | 13.93 | 13.77 | 11.07 | 16.91 | - | 2 |
| **5C** | CldU + IdU | R54 KO-1 | NT | 17.89 | 17.37 | 15.16 | 20.04 | <0.0001[§] | 2 |
| **5C** | CldU + IdU | R54/AP1 KO-2 | NT | 12.58 | 12.13 | 10.16 | 14.71 | <0.0001[§,†] | 2 |
| **5C** | CldU + IdU | R54/R54B KO-1 | NT | 15.14 | 14.70 | 12.06 | 17.73 | <0.0005[§] <0.0001[†] | 2 |

| | | | | | | | | | |
|---|---|---|---|---|---|---|---|---|---|
| **5C** | CldU + IdU | WT | Olaparib | 16.73 | 15.77 | 12.50 | 20.44 | <0.0001[$] | 2 |
| **5C** | CldU + IdU | R54 KO-1 | Olaparib | 22.86 | 23.43 | 19.73 | 26.65 | <0.0001[§] | 2 |
| **5C** | CldU + IdU | R54/AP1 KO-2 | Olaparib | 22.08 | 22.04 | 18.74 | 25.44 | <0.0001[§] 0.0160[†] 0.0140[¶] | 2 |
| **5C** | CldU + IdU | R54/R54B KO-1 | Olaparib | 20.97 | 21.19 | 17.83 | 24.1616 | <0.0001[§,†] | 2 |
| **S3B** | CldU | WT | NT | 9.87 | 9.93 | 8.94 | 10.91 | - | 2 |
| **S3B** | CldU | AP1 KO | NT | 9.88 | 9.83 | 8.35 | 11.15 | >0.9999[§] | 2 |
| **S3B** | CldU | R54 KO-1 | NT | 10.37 | 10.28 | 9.51 | 11.30 | 0.0035[§] | 2 |
| **S3B** | CldU | R54/AP1 KO-1 | NT | 9.83 | 9.77 | 8.92 | 10.66 | >0.9999[§] | 2 |
| **S3B** | CldU | R54/R54B KO-1 | NT | 9.35 | 9.30 | 6.97 | 11.78 | 0.2702[§] | 2 |
| **S3C** | CldU | WT | HU | 7.79 | 7.81 | 6.39 | 9.12 | - | 2 |
| **S3C** | CldU | AP1 KO | HU | 8.19 | 8.01 | 7.04 | 9.20 | 0.6179[§] | 2 |
| **S3C** | CldU | R54 KO-1 | HU | 8.54 | 8.42 | 7.34 | 9.71 | <0.0001[§] | 2 |
| **S3C** | CldU | R54/AP1 KO-1 | HU | 8.25 | 8.19 | 6.79 | 9.50 | 0.1752[§] | 2 |
| **S3C** | CldU | R54/R54B KO-1 | HU | 9.41 | 9.40 | 7.59 | 11.09 | <0.0001[§] | 2 |

[§] Compared to wild type (WT) HeLa cells.
[$] Compared to untreated (WT) HeLa cells (NT).
[†] Compared to RAD54 KO-1 cells.
[‡] Compared to RAD54/RAD54B KO-1 and RAD54/RAD54B KO-2 cells.
[¶] Compared to RAD54/RAD54B KO-1 cells.
[#] Compared to the corresponding non-treated (NT) cells.

**Table S2. Summary of comet assay analyses.** Means, medians, 25th and 75th percentiles, *p*-values and number of experiments (N), as shown in the graphs of the corresponding figures.

| Fig. | Genotype | Treatment | Mean (µm) | Median (µm) | 25th percentile | 75th percentile | *p* | N |
|---|---|---|---|---|---|---|---|---|
| **S5** | WT | NT | 59.78 | 56.76 | 44.68 | 73.43 | - | 3 |
| **S5** | R54 KO-1 | NT | 69.68 | 73.05 | 54.11 | 84.60 | <0.0001§ | 3 |
| **S5** | R54/AP1 KO-2 | NT | 66.81 | 67.73 | 46.96 | 83.54 | <0.0001§ 0.0418† | 2 |
| **S5** | R54/R54B KO-1 | NT | 64.43 | 60.01 | 46.40 | 80.08 | 0.0248§ 0.0001† | 2 |
| **S5** | WT | Olaparib | 74.76 | 76.77 | 60.65 | 90.64 | - | 3 |
| **S5** | R54 KO-1 | Olaparib | 74.37 | 78.69 | 53.54 | 92.89 | 0.9530§ | 3 |
| **S5** | R54/AP1 KO-2 | Olaparib | 97.12 | 97.52 | 86.22 | 110.19 | <0.0001§ <0.0001† 0.0563¶ | 2 |
| **S5** | R54/R54B KO-1 | Olaparib | 92.96 | 95.78 | 83.23 | 106.72 | <0.0001§ <0.0001† | 2 |

§ Compared to wild type (WT) HeLa cells.
† Compared to RAD54 KO-1 cells.
¶ Compared to RAD54/RAD54B KO-1 cells.

**Table S3. sgRNAs, primers and siRNAs used in this study.**

| sgRNAs | | |
|---|---|---|
| **Name** | **Target** | **Sequence (listed 5´- 3´)** |
| sgRNA A | *RAD54* (exon 8) | GCCTGGTGAAGAACTGGTAC |
| sgRNA B | *RAD54* (exon 8) | CGGAGGGAGGATCCAACCTC |
| sgRNA A | *RAD54B* (exon 6) | TAAGAACTGTTTCCCTCTTG |
| sgRNA B | *RAD54B* (exon 6) | TGGTGATTCTTATCTGGTCG |

| Primers | | | |
|---|---|---|---|
| **Primer** | **Target** | **Sequence (listed 5´- 3´)** | **Product Length (bp)** |
| P1 | *RAD54* (intron 7) | AGACTACCATCCCTGGGACA | 597 |
| P2 | *RAD54* (intron 8) | TGTTCCCTTTACACCTTTTCTGTTG | |
| P1 | *RAD54B* (intron 5) | TGAGAAGCTGTGAACATTGGC | 894 |
| P2 | *RAD54B* (intron 6) | CCACACTAGCAGTCGGTAAG | |

| siRNAs | |
|---|---|
| **Name** | **Target Sequence (listed 5´- 3´)** |
| non-depleting negative control | GATTCGAACGTGTCACGTCAA |
| *RAD51AP1* | AACCTCATATCTCTAATTGCA |
| *RAD54* | AAGCATTTATTCGAAGCATTT |
| *RAD54B* | ACCCAAGAAATTATAAATAAA |
